# Marginal Effects of Systemic CCR5 Blockade with Maraviroc on Oral Simian Immunodeficiency Virus Transmission to Infant Macaques

**DOI:** 10.1101/299206

**Authors:** Egidio Brocca-Cofano, Cuiling Xu, Katherine S. Wetzel, Mackenzie L. Cottrell, Benjamin B. Policicchio, Kevin D. Raehtz, Dongzhu Ma, Tammy Dunsmore, George S. Haret-Richter, Karam Musaitif, Brandon F. Keele, Angela D. Kashuba, Ronald G. Collman, Ivona Pandrea, Cristian Apetrei

## Abstract

Current approaches do not eliminate all HIV-1 maternal-to-infant transmissions (MTIT); new prevention paradigms might help avert new infections. We administered Maraviroc (MVC) to rhesus macaques (RMs) to block CCR5-mediated entry, followed by repeated oral exposure of a CCR5-dependent clone of simian immunodeficiency virus (SIV)mac251 (SIVmac766). MVC significantly blocked the CCR5 coreceptor in peripheral blood mononuclear cells and tissue cells. All control animals and 60% of MVC-treated infant RMs became infected by the 6th challenge, with no significant difference between the number of exposures (p=0.15). At the time of viral exposures, MVC plasma and tissue (including tonsil) concentrations were within the range seen in humans receiving MVC as a therapeutic. Both treated and control RMs were infected with only a single transmitted/founder variant, consistent with the dose of virus typical of HIV-1 infection. The uninfected RMs expressed the lowest levels of CCR5 on the CD4^+^ T cells. Ramp-up viremia was significantly delayed (p=0.05) in the MVC-treated RMs, yet peak and postpeak viral loads were similar in treated and control RMs. In conclusion, in spite of apparent effective CCR5 blockade in infant RMs, MVC had marginal impact on acquisition and only a minimal impact on post infection delay of viremia following oral SIV infection. Newly developed, more effective CCR5 blockers may have a more dramatic impact on oral SIV transmission than MVC.

**Importance:** We have previously suggested that the very low levels of simian immunodeficiency virus (SIV) maternal-to-infant transmissions (MTIT) in African nonhuman primates that are natural hosts of SIVs are due to a low availability of target cells (CCR5^+^ CD4^+^ T cells) in the oral mucosa of the infants, rather than maternal and milk factors. To confirm this new MTIT paradigm, we performed a proof of concept study, in which we therapeutically blocked CCR5 with maraviroc (MVC) and orally exposed MVC treated and naïve infant rhesus macaques to SIV. MVC had only a marginal effect on oral SIV transmission. However, the observation that the infant RMs that remained uninfected at the completion of the study, after 6 repeated viral challenges, had the lowest CCR5 expression on the CD4^+^ T cells prior to the MVC treatment, appear to confirm our hypothesis, also suggesting that the partial effect of MVC is due to a limited efficacy of the drug. Newly, more effective CCR5 inhibitors may have a better effect in preventing SIV and HIV transmission.

## Introduction

Despite enormous success in preventing mother-to-infant-transmission (MTIT), recently, the World Health Organization (WHO) has intensified international efforts to significantly reduce or eliminate infection of infants. In 2013, UNAIDS reported that approximately 210,000 infants worldwide become HIV-infected annually (1). More than 90% of these HIV-1 infections occur in sub-Saharan Africa. MTIT can occur *in utero*, directly by hematogenous transplacental spread or by infection of the amniotic membranes and fluid (2); during the delivery, by contact of the infant with maternal blood and cervicovaginal secretions (3, 4); or postnatally, through breastfeeding (5, 6). This later mode of transmission accounts for most MTIT cases and is difficult to prevent, because its mechanisms are not completely understood. Differently from HIV vaginal or rectal transmission, in which the virus-host interactions are intensively studied at the portal of entry (7), little emphasis has been placed on the role of infant mucosa in HIV breastfeeding transmission. This paucity of information is mainly due to the inherent limitations of sampling human infants. Further challenges to studying infant oral transmission include the long duration of exposure from breast milk and dramatic age-related changes in the infant mucosa during that time. In addition, most HIV-infected women are receiving some form of antiretroviral therapy (ART) or peripartum prophylaxis (8), which reduces MTIT but makes it more difficult to study break-through infections. As such, MTIT studies have focused almost exclusively on maternal virological and immunologic factors (9-11) and on immune effectors present in breast milk (12-16). High HIV-1 maternal plasma viral load (VLs) and low CD4^+^ T cell counts in women that breastfeed are correlated with increased HIV breastfeeding transmission (17, 18), but these correlations are not always substantiated, as mothers with low VLs can also transmit HIV by milk (17, 18). Conversely, 63% of the infants breastfed by mothers with <200 CD4^+^ T cells/µL and >10^5^ vRNA copies/ml remain uninfected (19). Furthermore, the correlation between milk viral shedding and plasma VL is weak and substantial discrepancies exist, with some women having low VLs in milk but high VLs in plasma, and *vice versa* (19). The rates of breastfeeding transmission are also correlated with the duration of lactation rather than the absolute CD4^+^ T cell count (20). These data highlight the complex and dynamic process of infant oral transmission.

Breastfeeding transmission studies in macaques have also only focused on maternal and milk factors (13, 16, 21, 22). Neither maternal plasma VLs nor CD4^+^ T cells clearly predict breastfeeding transmission in macaques, with only 20% of acutely-infected dams successfully transmitting infection. Importantly, over 50% of SIV breastfeeding transmissions occurred 9 months postdam infection, when the offspring are older, highlighting an age-related susceptibility to SIV infection, with higher doses of virus needed to infect younger RMs (23). Finally, it has been reported that occult peripartum/postpartum SHIV infection that may occur early may go undetected until later, suggesting that maturation of the immune system and generation of target cells in the infant are needed to support virus replication (21).

Our previous work in African nonhuman primates (NHPs) that are natural hosts of SIVs demonstrated that in these species MTIT of SIV is virtually nonexistent (<5%) (24-26) and below the level targeted by the WHO for “virtual elimination” of HIV-1 MTIT in humans (27). The low levels of MTIT in natural hosts contrast with massive offspring exposure to SIV both *in utero* and through breastfeeding (25) due to the high SIV prevalence in the wild (>80%) and high levels of acute and chronic viral replication in dams (25, 26). In African green monkeys (AGMs) and mandrills, resistance to SIV breastfeeding transmission is strongly associated with low levels of SIV target cells at the mucosal sites of the offspring (24). Furthermore, AGM-susceptibility to experimental SIV mucosal transmission is proportional to the availability of CD4^+^ T cells expressing the SIV coreceptor CCR5^+^ at the mucosal sites (28, 29).

Based on these observations, we hypothesized that the levels of target cells (CCR5^+^ CD4^+^ T cells) at the oral mucosa of breastfed infants may drive the efficacy of HIV/SIV transmission through breastfeeding and that the CCR5 blockade could represent a new potential therapeutic strategy to prevent HIV/SIV breastfeeding transmission. We tested this hypothesis in an infant RM model of HIV breastfeeding transmission (16), in which we administered Maraviroc (MVC) to block oral SIVmac transmission. MVC was shown to effectively block CCR5 expression in mucosal CD4^+^ T cells, and prevent SIV transmission upon topic administration (30), but systemic CCR5 blockade to prevent oral HIV/SIV transmission has never been performed. MVC has low toxicity (31) and high penetrability to the mucosal sites and is available for oral administration, thus being suitable for the use in infants. As such, we reasoned that demonstrating MVC efficacy in blocking oral HIV transmission may lead to an efficient way to prevent HIV breastfeeding transmission. We report here that, while systemic MVC administration to infant RMs was well tolerated and efficiently blocked CCR5 in peripheral blood and at mucosal sites, it had a minimal impact on viral acquisition and only marginally impacted post infection delay of viremia. The infant RMs that remained uninfected at the completion of the study had the lowest CCR5 expression on the CD4^+^T cells prior to the MVC treatment, confirming our hypothesis that the availability of target cells may drive the efficacy of SIV/HIV breastfeeding transmission, also suggesting that the partial effect of MVC is due to a limited efficacy. Newly, more effective CCR5 inhibitors may have a better effect in preventing SIV and HIV transmission.

## Results

### Study design

To investigate whether or not blockade of the mucosal target cells can prevent/reduce HIV/SIV oral transmission, five infant RMs were administered MVC at a total daily dose of 300 mg/kg bid (150 mg/kg given twice daily), by mouth with food, for up to 4 months. One month after MVC initiation, the treated infants, together with four uninfected controls, received 10,000 IU of SIVmac766XII (a synthetic swarm of the transmitted/founder SIVmac766 clone) (Figure 1) (32) via oral, atraumatic administration. Viral challenges were repeated every two weeks until all the controls became SIV-infected (after the 6^th^ challenge).

**Figure 1.**
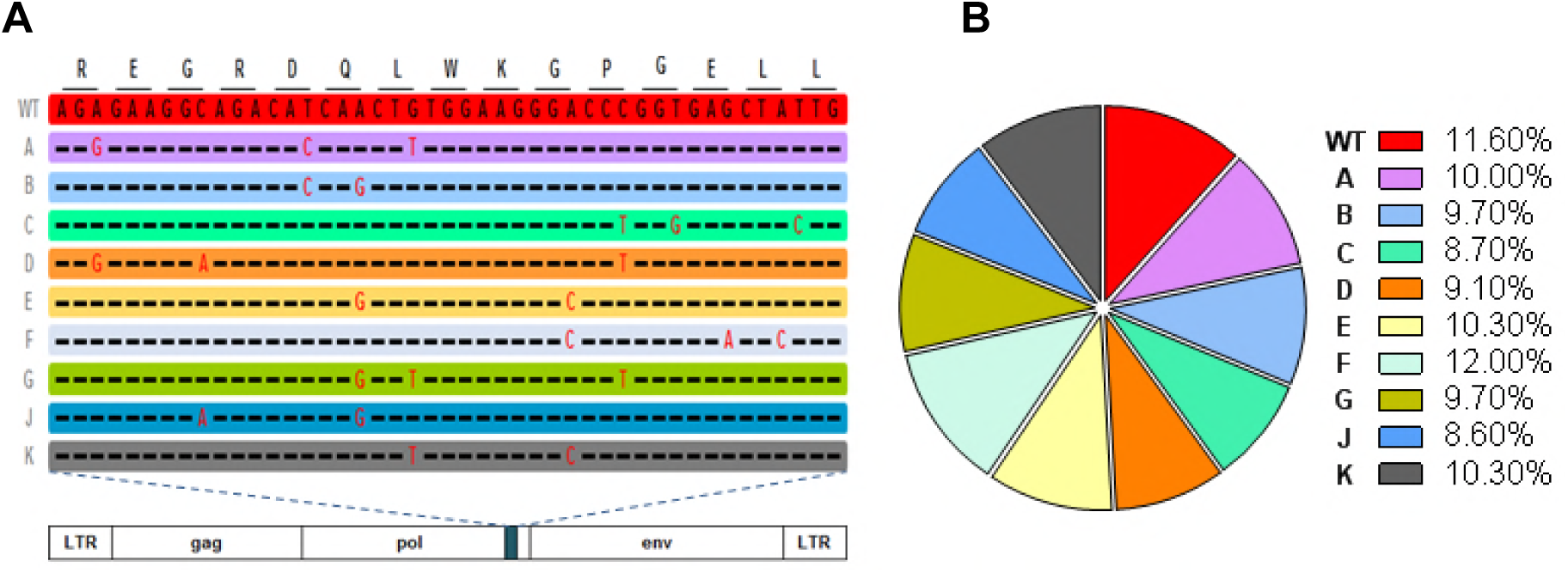
Alterations in the SIVmac766 clone that allow for discriminating the number of unique T/F variants. SIVmac766XII is an infection stock is composed of eleven distinct viral clones differing from wild-type virus by 3 synonymous mutations within the integrase gene (A). The entire remaining genome is identical between clones. The proportion of each variant in the viral stock was determined by RT-SGA with 334 sequences examined (B). All mutations and the pie chart are color coded for each of the twelve clones within the synthetic swarm.

At the time of viral challenges, MVC was dosed in circulation in all the MVC-treated infant RMs. Due to the nature of the study, which involved repeated oral challenges, we did not collect oral or tonsil biopsies to dose the MCV at the site of virus exposure, to avoid increasing the risk of SIV transmission. However, we assessed the MVC concentration in tissues (including tonsils) in two additional MVC-treated SIV-unchallenged infant RMs, which were followed in the same conditions as the infants in the study group.

Blood (1.5 ml) was collected into EDTA-Cell Preparation Tubes (CPTs) from all the infant RMs receiving MVC at the time of challenge, to monitor coreceptor occupancy (33) and measure the plasma concentrations of MVC. Blood was then collected every three days to detect the SIV infection. Once an animal was diagnosed as SIV-infected, frequent blood samples were collected to monitor the acute and early chronic infection (10, 17, 24, 31, 38, 45, 59 day postinfection, dpi). Superficial lymph nodes (LNs), tonsils and gut biopsies were collected only from the RMs in the MVC-treated control group.

### Orally administered MVC is well tolerated by infant RMs

Throughout the MVC treatment (up to 101 days), all infant RMs receiving MVC were closely monitored for clinical or biological signs suggestive of side or adverse effects of the MVC. No such signs being observed, we concluded that oral administration of MVC was safe and well tolerated by infant RMs.

### Pharmacokinetics (PK) of MVC in plasma and tissues

The PK profile of MCV was evaluated in all the infant RMs from the study group by measuring the MVC plasma concentrations 4 hours after the morning administration, when we expected drug levels to be maximal and when viral challenges were performed. Additional testing of the MVC plasma concentrations was performed at 2, 3 and 7 days postviral challenge, just before the morning administration of the MVC, when we expected the plasma concentrations to be minimal (Figure 2). The medians and ranges plasma MVC concentrations at the time of each of the 6 virus challenges were respectively of 410 (77-1040), 886 (29-1910), 115 (23-267), 1960 (815-3720), 1435 (5-5340) and 64 (27-1520) ng/ml, respectively (Figure 2). In the unchallenged MVC-treated controls, plasma MVC concentrations were in the same range: 248 and 261 ng/ml (Figure 3A). These levels are similar to the range seen in humans receiving a single 300 mg dose of MVC (618-888 ng/ml) (34, 35). The medians and ranges of the MVC concentrations in plasma just prior to the morning dose (the minimal coverage concentration) were of 59 ((25-271), 46 ((15-214), 28 ((13-144),11 (5-21), 33 (21-62) and 25 (19-44) ng/ml at 3 days post-challenges, demonstrating a steady and measurable MVC trough levels. In the MVC-treated controls, the minimal concentrations of MVC were of 207 and 33 ng/mL (Figure 3A). Overrall, these levels were slightly lower than those measured at the same interval post-MVC administration in humans receiving a single 300 mg dose (34-43 ng/mL) (34, 35).

**Figure 2.**
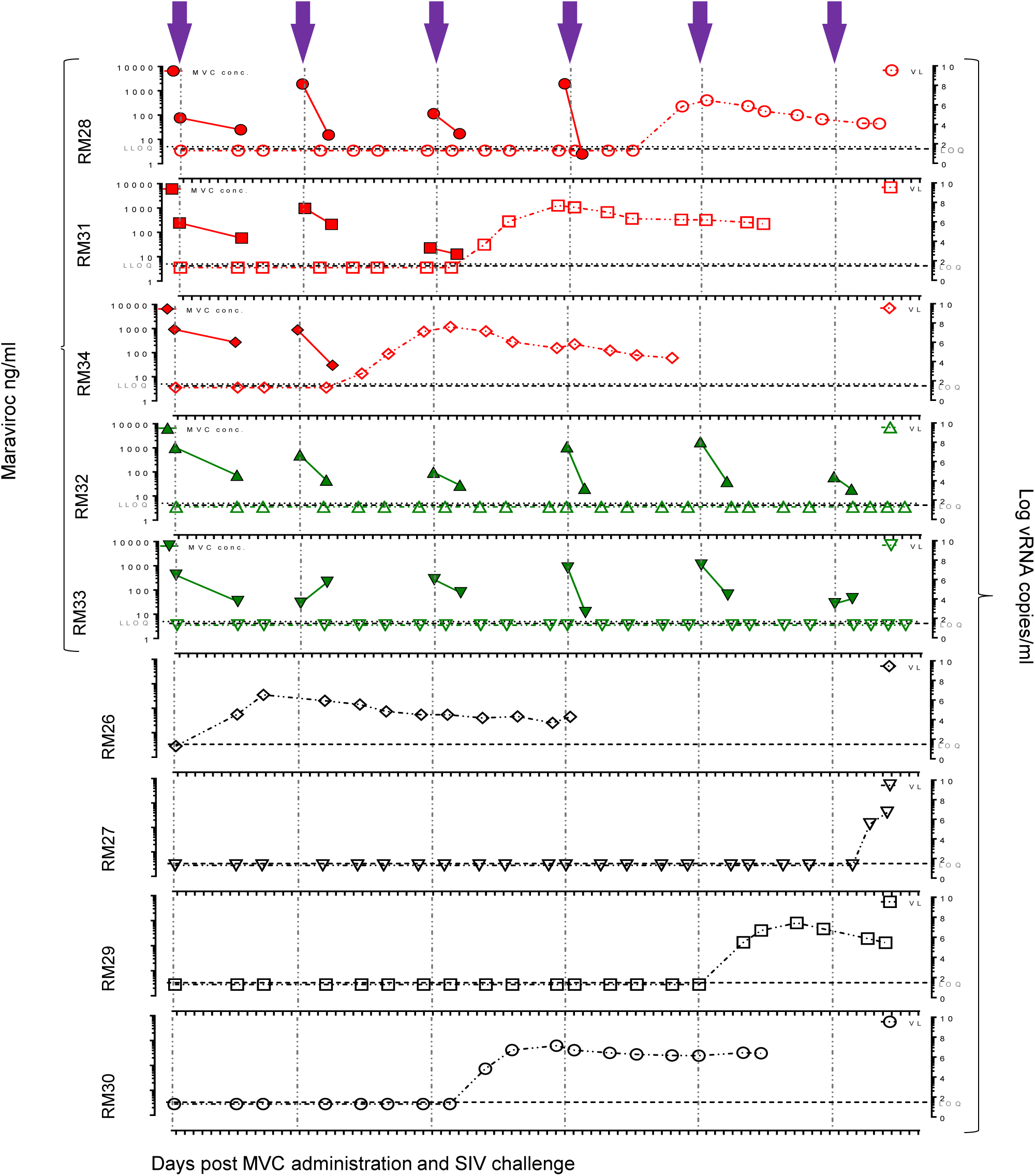
Comparative assessment of MVC pharmacokinetics and plasma VLs at the time of and after the SIVmac766XII challenge. MVC concentrations in plasma at 4 hours (maximum concentration) and 16 h (minimum concentration) after systemic administration of 150 mg/kg of MVC; Plasma VLs are shown at the corresponding time points of treatment and viral challenge for infant RMs in the MVC-treated group and untreated controls. Closed symbols represent MVC concentration, open symbols illustrate the viral loads. MCV is expressed as ng/mL of plasma, the viral load is expressed as logarithms of the numbers of viral RNA copies per ml of plasma. Gray dotted line show the lower limit of quantification (LLOQ, 5 ng/ml) of the bioanalytical LC-MS/MS method; short dashed line shows the limit of viral load quantification (LOQ, 30 copies per ml). Violet arrows illustrate the virus challenge.

**Figure 3.**
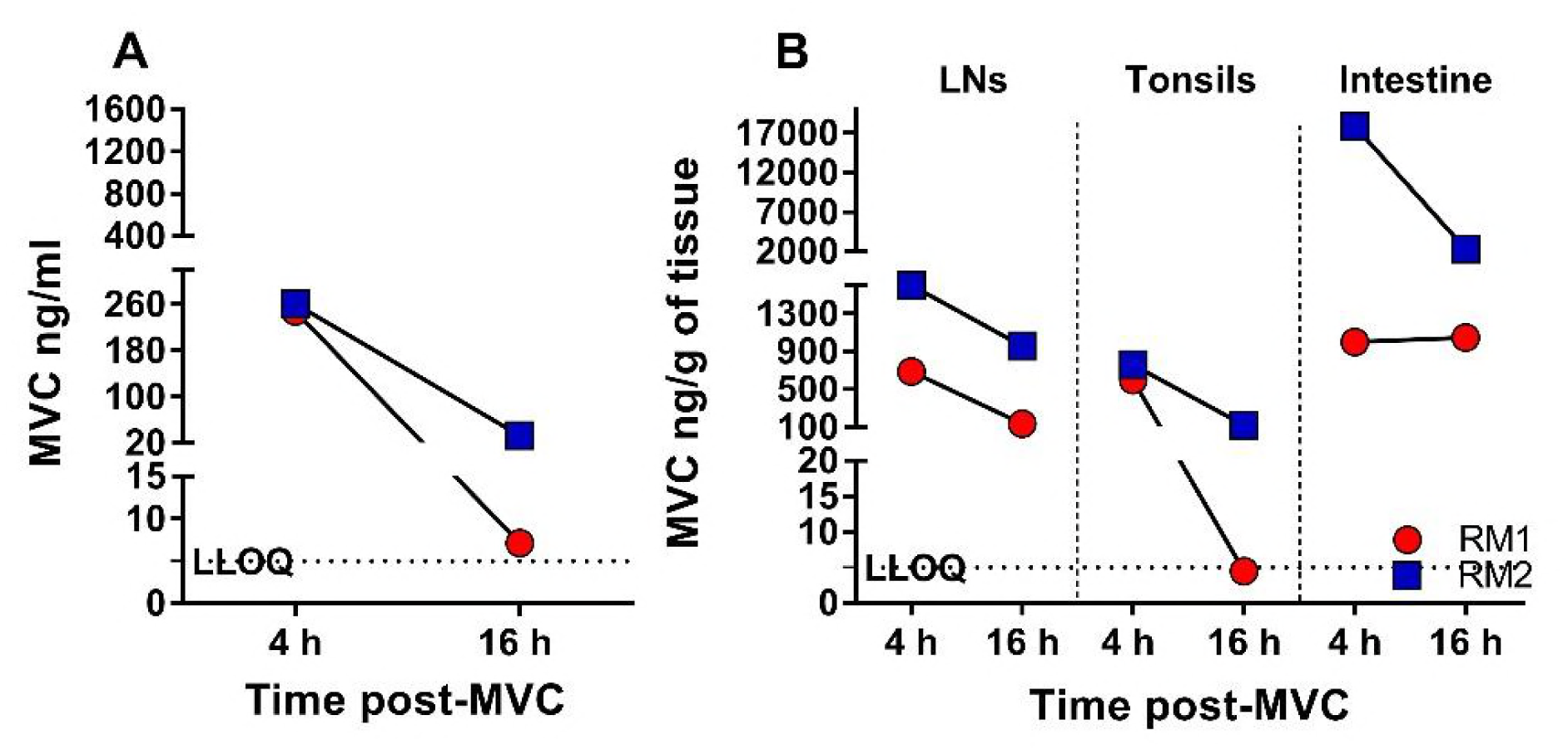
Pharmacokinetic analysis of MVC concentration in plasma and tissues in two infant RM from the MVC-treated control group. (a) MVC concentrations in plasma at 4 hours (maximum concentration) and 16 h (minimum concentration) after systemic administration of 150 mg/kg of MVC; MCV concentration is expressed as ng/mL of plasma. (b) MVC concentrations in LNs, tonsils and intestine at 4 hours and 16 h after systemic administration; MCV concentration is expressed as ng/g tissue. Red circles represent RM1, blue squares represent RM2. Black dotted line show the lower limit of quantification (LLOQ, 5 ng/ml) of the bioanalytical LC-MS/MS method

At 4 hours after the drug administration, the MVC concentrations in the tissues collected from the MVC-treated controls were 689 and 1597 ng/g in the LNs, 597 and 759 ng/g in the tonsils, and 998 and 17,869 ng/g in the gut (Figure 3B). The MVC concentrations in tissues immediately prior to the morning dose were 136 and 958 ng/g in the LNs, 5 and 122 ng/g in tonsils, and 1,046 and 2,322 ng/mL in the gut (Figure 3B). In only two of the collected samples (plasma from RM28 2 days postchallenge 4 and tonsil from RM1) MVC concentrations were below the 5 ng/ml limit of quantification (BLQ) of the method used (Figures 2 and 3). We imputed a numerical value for these samples (5 ng/ml) because it was within 20% of the low limit of quantification (LLOQ) (36). Interestingly, in RM28, the MRV concentration below the limit of quantification was followed by SIV infection (Figure 2).

Taken together, these data demonstrate that an oral MVC dose of 150 mg/kg bid given to infant RMs 4 hours prior to the viral challenge, approximated plasma MVC concentration in humans; that the tissue concentrations of MVC were similar to those observed in humans (37) and high enough to block CCR5, and that the minimal concentrations of MVC were generally sufficient to compete with the virus for CCR5 coreceptor occupancy, albeit the concentrations of MVC decreased dramatically prior to the daily administration, in some instances, below 5 ng/ml.

### Orally administered MVC effectively blocks CCR5 expression on the surface of CD4^+^ T cells

To investigate whether or not CCR5 blockade with MVC impacts oral SIV transmission to infant RMs, we first determined the therapeutic impact of MVC by measuring the CCR5 receptor occupancy in blood, LNs, tonsil and gut. This test monitors the levels of internalization of CCR5 receptors on the surface of CD4^+^ T cells following *ex vivo* MIP-1β exposure, which are indicative of the level of receptor occupancy. Complete prevention of CCR5 internalization indicates complete coreceptor occupancy.

Close monitoring of CCR5 occupancy on the surface of circulating CD4^+^ and CD8^+^ T cells (Figure 4) identified significant differences between MVC-treated and untreated groups before the first viral challenge (CD4^+^ T cell CCR5 occupancy p=0.0159; CD8^+^ T cell CCR5 occupancy p=0.0317), before the second viral challenge (CD4^+^ T cell CCR5 occupancy p=0.0317; CD8^+^ T cell CCR5 occupancy p=0.0159), and before the third viral challenge (CD4^+^ T cell CCR5 occupancy p=0.0286; CD8^+^ T cell CCR5 occupancy p=0.0159) (Figure 4A and B). For the remaining 3 challenges, statistical analyses could not be performed because the number of uninfected RMs were too low.

**Figure 4.**
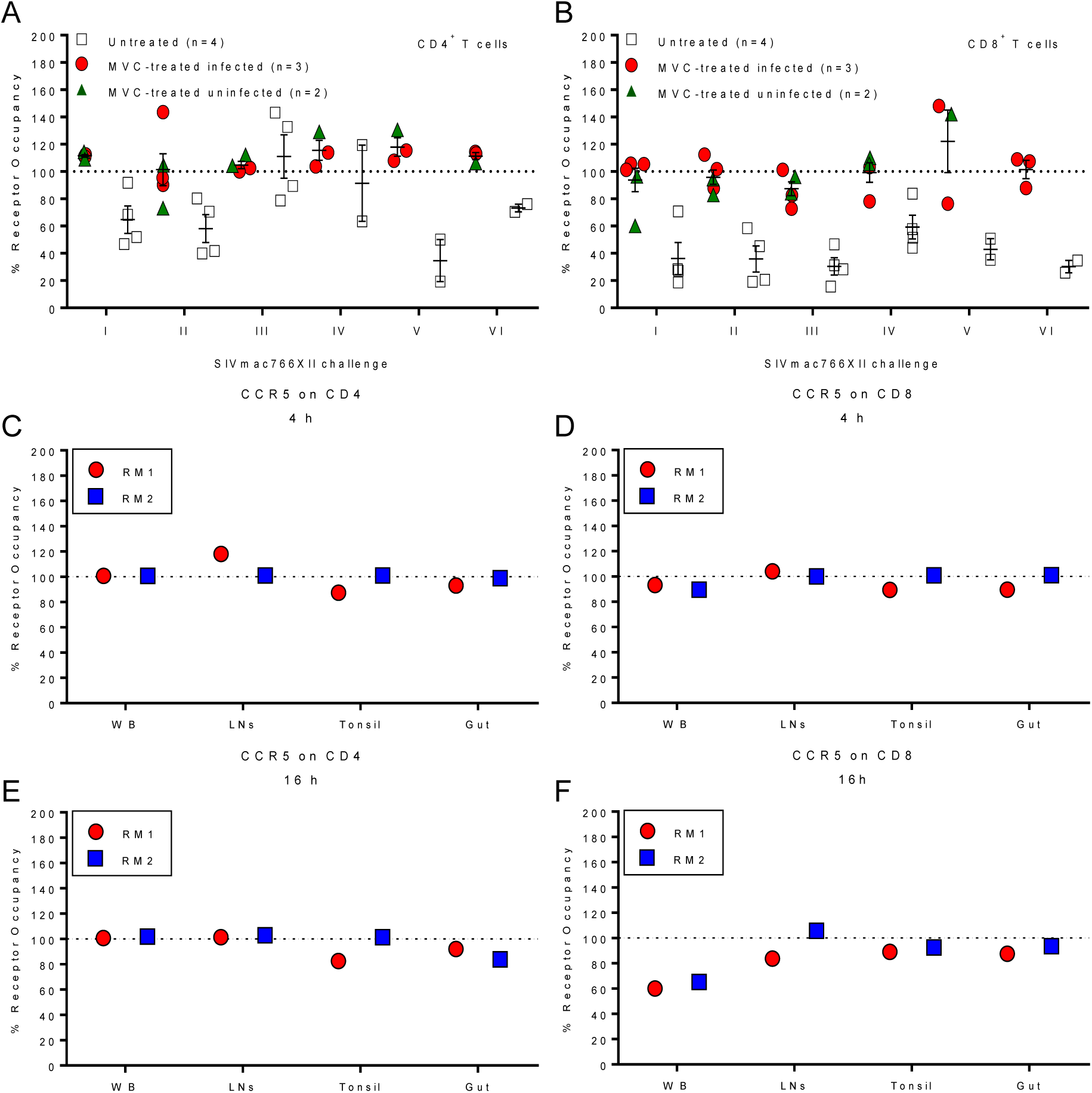
CCR5 receptor occupancy on CD4^+^ and CD8^+^ T cells from blood, lymph nodes, tonsils and intestine from two infant RMs. Percentage of CCR5 receptor occupancy on circulating CD4^+^ T cells (a) and CD8^+^ T cells (b) from infant RMs included in the study group at the time of SIVmac challenge. White open squares represent untreated infant RMs from the control group, red circles represents MVC-treated infant RMs and green triangle represent MVC-treated uninfected infant RMs. Data are presented as individual values with the group means (long solid lines) and standard errors of the means (short solid lines). Mann-Whitney test was used to calculate the exact p value. (c-f) Coreceptor occupancy in blood and tissues of the infant RMs from the MVC-treated control group. Percentage of receptor occupancy on CD4^+^ and (e) CD8^+^ T cells 4 h after the MVC administration (maximum concentration). (d) Percentage of receptor occupancy on CD4+ and (f) CD8+ T cells 16 h after MVC administration (maximum concentration). Red circles represent infant RM1, blue squares represent infant RM2.

In the MVC-treated controls, MVC blocked efficiently CCR5 on CD4^+^ T cells in all tissue samples analyzed (Figure 4C and D). In the gut, CCR5 blockade was not complete, even though blocking efficiency was high with average levels of 96% when the MVC concentration was expected to be high (Figure 4C) and 88% when the MVC concentration was expected to be low (Figure 4D). Similarly, MVC partially blocked the CCR5 expression on the CD8^+^ T cells (Figure 4E and F), with an average CCR5 occupancy of 91% (when the MVC concentration was expected to be high) and 63% (when the MVC’s concentration was expected to be low) in whole blood; blockade was of 102% and 95% in the LNs; 95% and 91% in the tonsil and 95% and 91%, respectively in the gut (Figure 4).

### Systemic MVC administration only marginally impacted oral SIVmac transmission to infant RMs

The main goal of this study was to investigate whether or not CCR5 blockade with MVC impacts oral SIV transmission to infant RMs. MVC-treated and control infant RMs were repeatedly challenged with 10,000 IU of SIVmac766XII, orally, in an atraumatic fashion, until all 4 RM controls became infected (6 challenges). At the end of the challenge experiments, 3/5 (60%) of the MVC-treated infant RMs were also SIV-infected, while 2/5 infant RMs remained uninfected, in spite of being challenged 6 times under the same conditions (Figure 2). However, the levels of protection in the MVC-treated RMs were not significant (p=0.15). We conclude that systemic MVC administration does confer a significant protection of the infant RMs against oral SIVmac challenge. This conclusion is also supported by the observation that the number of exposures necessary to infect the infant RMs in the two groups were similar, control infant RMs becoming infected after 1, 3, 5 and 6 SIVmac766XI oral challenges, respectively, and the MVC-treated infant RMs becoming infected after 2, 3 and 4 inoculations.

We next sought to correlate the efficacy of the SIVmac766XII transmission (estimated based on the number of virus challenges) with the availability of CCR5^+^ CD4^+^ T target cells. This analysis was prompted by our previous correlative studies in natural hosts of SIVs that found strong correlations between the target cell availability at mucosal sites and the efficacy of mucosal (intrarectal, intravaginal and oral) transmission (24, 28). We assessed CCR5 expression on circulating CD4^+^ T cells of the infant RMs prior to the MVC treatment and correlated it to the number of viral exposures prior to infection. In a conservative approach, we listed the uninfected RMs as infected at the seventh challenge. These two variables were very strongly correlated (p=0.0036) (Figure 5), confirming our hypothesis and strongly supporting the paradigm that target cell availability determines susceptibility to infection in natural hosts of SIVs.

**Figure 5.**
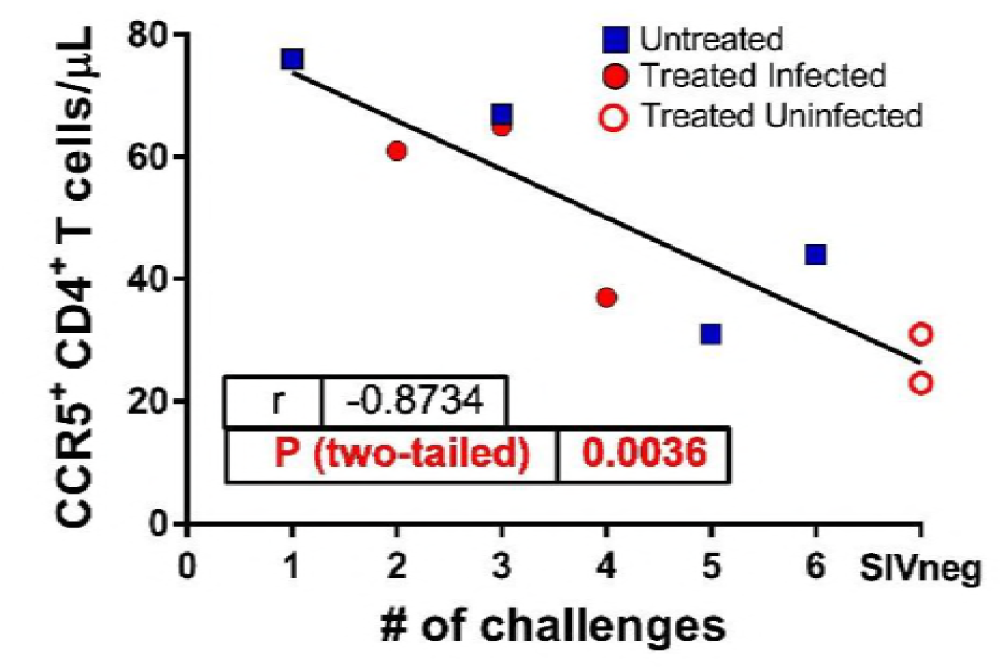
Correlation between the levels of CCR5 expression on the peripheral CD4^+^ T cells and the number of viral challenges required for infecting MVC-treated and untreated RMs. The two MVC-treated, SIV uninfected RMs are illustrated as open circles.

The SIVmac766XII stock consists of a swarm of 12 viral variants equally represented and phenotypically matched allowing for variant enumeration (Figure 1) (38), therefore the number of transmitted viral variants in the MVC-treated group and the untreated controls were determined. The number of transmitted/founder lineages, did not identify any difference in the number of transmitted variants between the two groups, with each animal being infected with only one of the 12 possible variants. This result suggests that the infant RMs were not overexposed to virus, which could have offset the protective effect of MVC.

### SIVmac766XII uses CCR5 and GPR15 to enter transfected target cells

To understand why the MVC administration only marginally impacted oral SIV transmission in infant RMs, we first investigated the coreceptor usage of SIVmac766XII. Several SIVsmm strains from sooty mangabeys were reported to use CXCR6 (39, 40), which, if true for SIVmac, could have resulted in a more promiscuous coreceptor use and the inefficacy of the CCR5 blockade. First, we assessed SIVmac766XII coreceptor usage in a CF2th-Luc reporter cells system, and documented robust viral entry through both RM CCR5 and RM GPR15 (Figure 6), but only minimal entry through RM CXCR6, and no virus entry through RM CXCR4, in agreement with previous studies of coreceptor usage of the SIVmac strains (39). As controls, other SIVsmm strains showed a robust entry through sooty mangabey CXCR6 (SM CXCR6, black bar, Figure 6) as previously reported (40).

**Figure 6.**
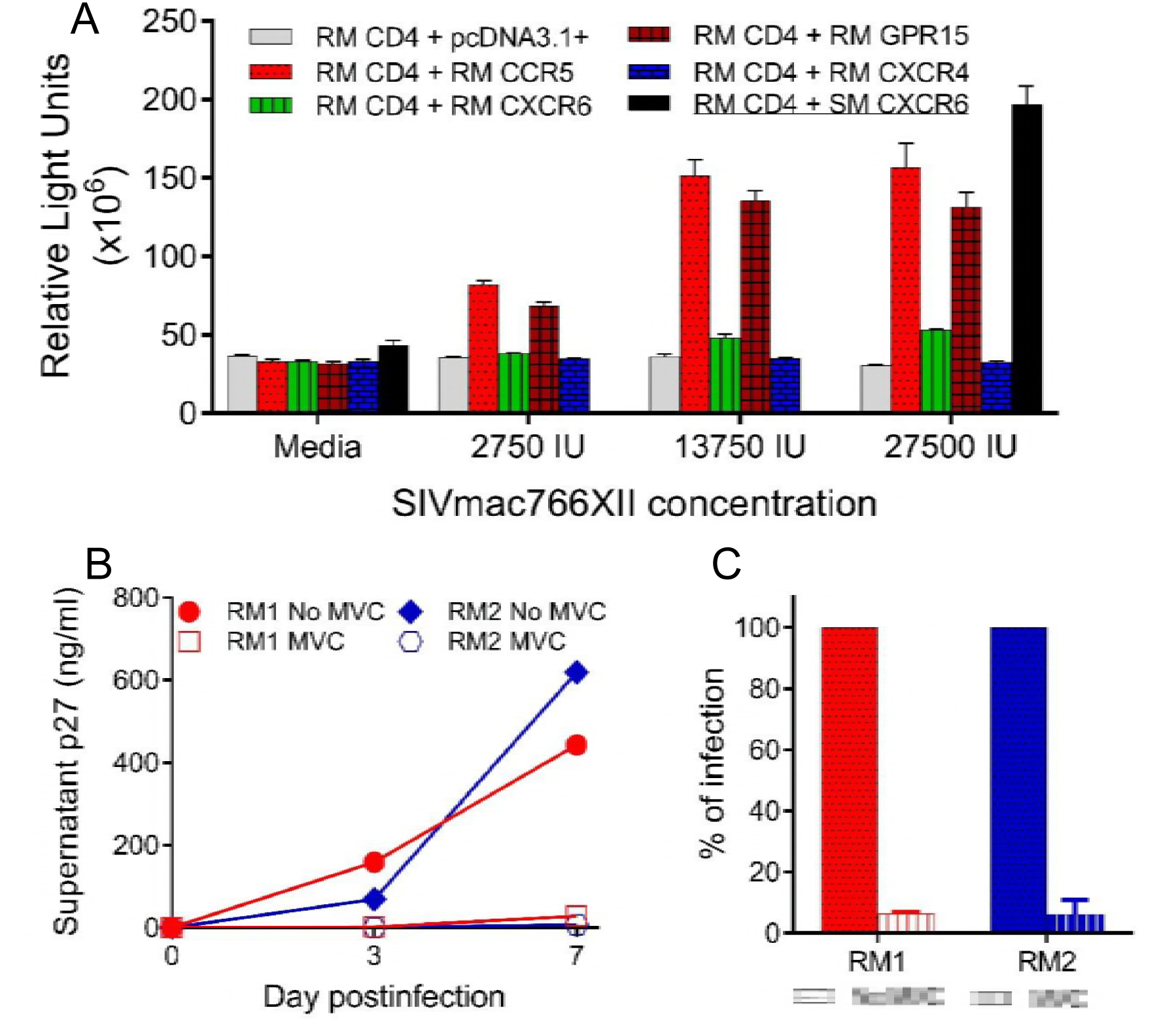
SIVmac766XII use of RM coreceptors. (A) CF2th-Luc cells that contain a Tat-driven luciferase reporter were transfected with expression plasmids containing RM CD4 and coreceptor. Cells were infected 48 hours later with SIVmac766XII (2750IU, 13750IU and 27500IU) and entry was quantified 72 hours later by measuring luciferase production as relative light units (RLU). Infections were carried out in triplicate and bars represent means and standard error of the mean (sme) values. Sooty mangabey CXCR6, which is a functional coreceptor, was included at the highest inoculum for comparison (SM CXCR6) black bar. (B) SIVmac766XII infectivity on PBMCs. PBMC from two RM were stimulated for three days with ConA and IL-2, then pretreated for one hour with maraviroc (MVC; 15 uM) or with vehicle alone (No Drug), and infected with SIVmac766XII (550IU). (C) SIVmac766XII infectivity on PBMCs. Infections were carried out in duplicate, and infection was measured by p27 production in the supernatant. Each line indicates one infection condition per animal, and data represents the mean and standard error of the mean values.

### CCR5 is the main coreceptor used by SIVmac766XII to infect primary RM PBMCs

We further assessed the SIVmac766XII coreceptor usage during infection of RM primary lymphocytes. PBMC from two different RMs were infected with SIVmac766XII in the presence or absence of MVC, and virus replication was monitored by measuring p27 Gag antigen in the supernatant. As shown in Figure 6B, MVC blocking of CCR5 virtually abolished infection of primary lymphocytes, with 94-99% reduction of replication at 7 dpi (Figure 6C). We therefore concluded that CCR5 is the main coreceptor for SIVmac766XII in primary PBMC, despite *in vitro* use of both RM CCR5 and RM GPR15 in transfected cells. This finding is concordant with previous results that SIVmac is dependent on CCR5 for primary lymphocyte infection (40, 41). As such, our results showed that SIVmac766XII is an appropriate viral strain to model oral transmission of HIV-1.

### Postinfection effects of the MVR treatment

We further analyzed the impact of the MVC treatments on the natural history of SIV infection in infant RMs. In these studies we included the three MVC-treated the four untreated infant RMs that became infected with SIVmac766XII.

### MVC treatment delayed ramp-up viremia

A significant delay of the ramp-up VLs was observed in the MVC-treated infants (p=0.05) (Figure 7). In addition to the delay in ramp-up dynamics, the peak VL for MVC-treated animals was reached at 21 days postinfection (dpi) compared to 18 dpi peaks of the control RMs. However, the MVC effects on timing and magnitude of pVL postpeak resolution and later in infection were not significantly different between the two groups (Figure 7).

**Figure 7.**
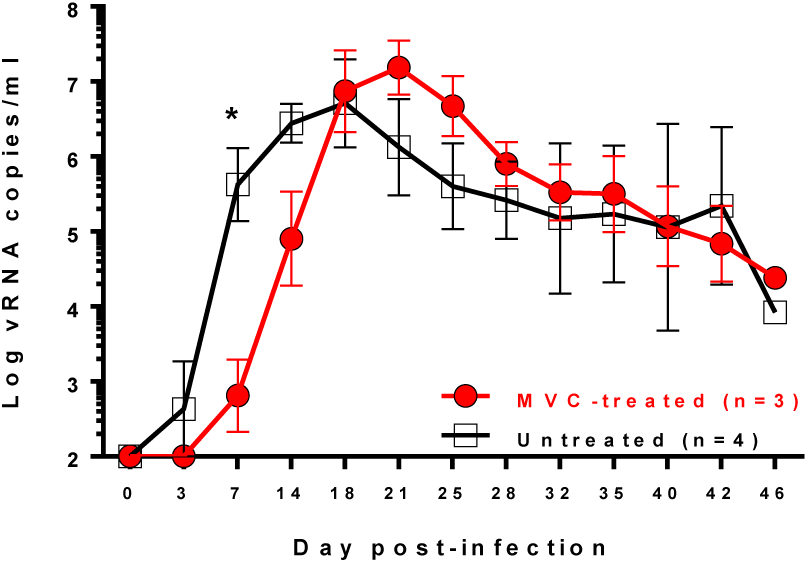
Changes in the viral loads of the SIVmac-infected RMs treated with MVC compared with untreated controls. Plasma vRNA loads (copies/ml, expressed in logarithmic format) in MVC-treated and untreated group. Data are geometrical means, with the bars representing standard error of the mean. MVC-treated infant RMs are representing as red circles; infant RM controls are showed as open black squares. Mann-Whitney test was used to calculate the exact p value (p= 0.05).

### MVC treatment did not alter the dynamics of the peripheral CD4^+^ and CD8^+^ T cell populations or subsets in SIV-infected infant RMs

In humans, MVC treatment does not significantly impact CD4^+^ and CD8^+^ T cell populations (42). The peripheral CD4^+^ (Figure 8A) and CD8^+^ (Figure 8B) T cell counts were compared throughout the follow-up between MVC-treated and untreated RMs, and no significant difference was observed between the two groups (Figure 8). Peripheral CD4^+^ T cell depletion was transient, the CD4^+^ T cell counts being partially restored by 24 dpi in both groups and declining slowly during the follow-up (Figure 8A).

**Figure 8.**
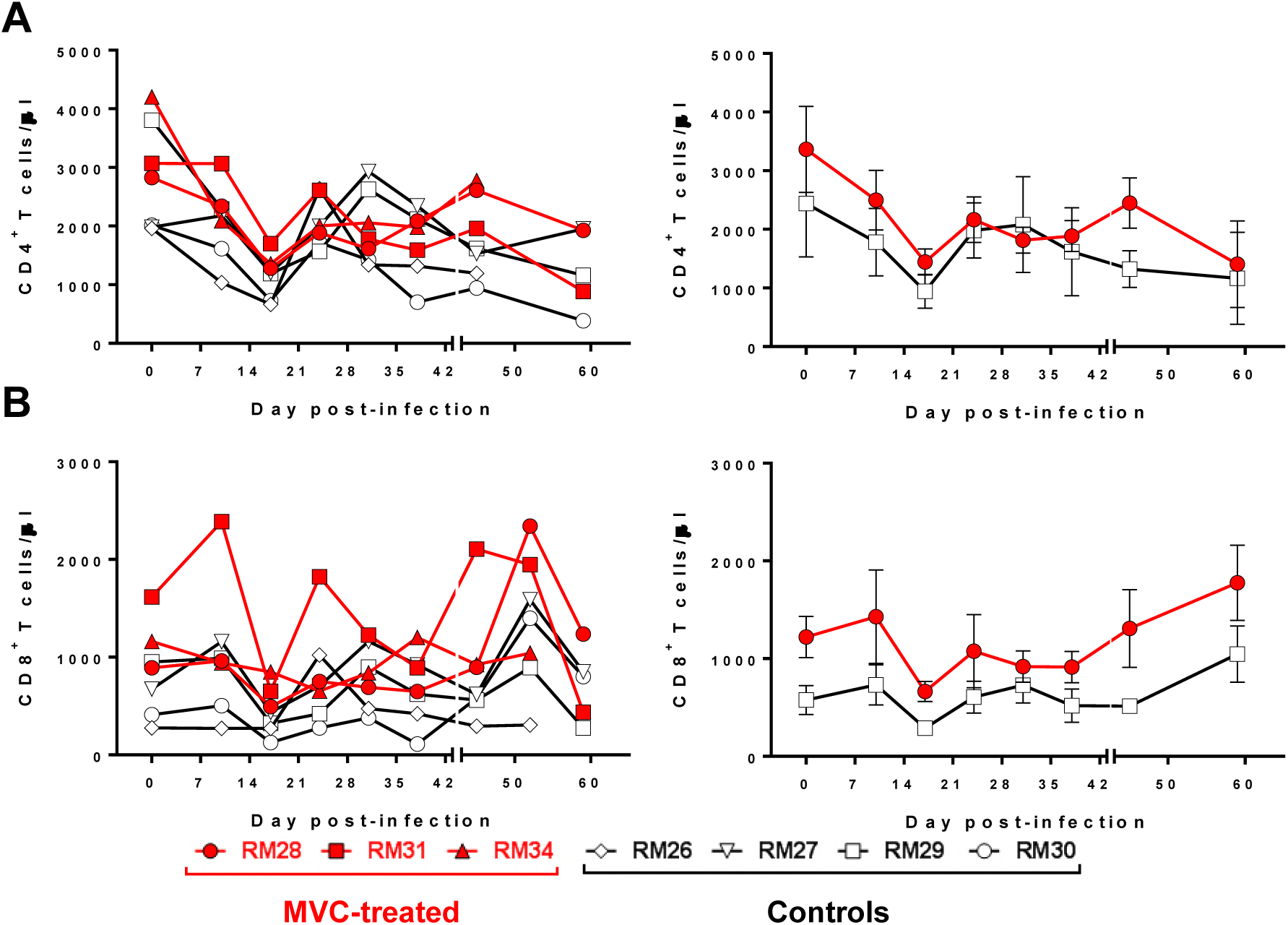
Longitudinal analysis of absolute CD4^+^ and CD8^+^ T cell counts in blood from the SIV-infected infant RMs. (A) Changes in the CD4^+^ T cells; (B) Changes in the CD8^+^ T cells; Left panels illustrate individual animals; right panels averages. Red symbols illustrate MVC-treated animals. Open black symbols illustrated untreated controls. Vertical bars in the right panels are the standard errors of the means.

We next monitored the impact of MVC treatment on the memory subsets of CD4^+^ and CD8^+^ T cells prompted by a recent report that CCR5 blockade *in vivo* might affect the trafficking of memory T cells expressing CCR5 to the site of the cognate antigen, preventing their proper stimulation, acquirement of effector functions and antiviral activity (43). However, comparison between MVC treated and untreated infant RMs throughout the follow-up did not reveal any significant difference in the peripheral naïve (CD28^+^ CD95^neg^), central memory (CD28^+^ CD95^+^) and effector memory (CD28^neg^ CD95^+^) subsets of CD4^+^ or CD8^+^ T cells (data not shown). Our data indicate that MVC treatment had no discernible impact on the major T cell populations and subsets in the SIV-infected infant RMs.

### MVC administration did not impact the levels of circulating CD4^+^ and CD8^+^ expressing CCR5 in SIV-infected infant RMs

CCR5 expression on the surface of CD4^+^ T cells is highly variable, depending on CCR5 polymorphisms and expression of its chemokine ligand (44, 45), leading to variations in HIV target cell availability, that impact virus entry, susceptibility to infection (46), and the therapeutic efficacy of CCR5 inhibitors (47). We therefore monitored CCR5 expression on both CD4^+^ and CD8^+^ T cells throughout the follow-up (Figure 9) and report that they were similar between the two groups, being increased during the first weeks of treatment (Figure 9) and returning to preinfection levels by 28 dpi. The CD4^+^ T cells expressing CCR5 gradually declined during the follow-up (Figure 9A and B) likely as a result of the direct virus targeting of the CD4^+^ T cells expressing CCR5.

**Figure 9.**
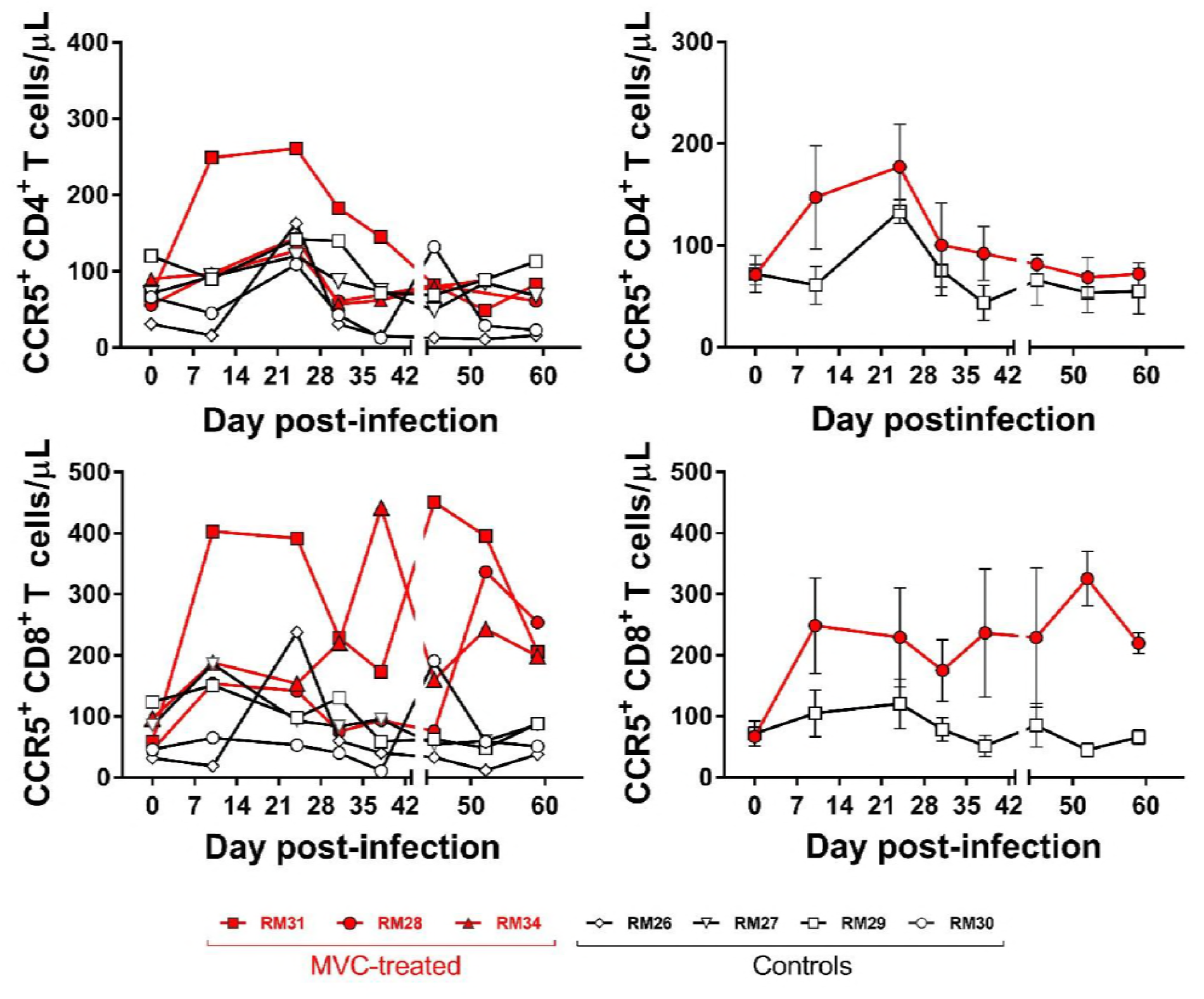
Longitudinal analysis of absolute counts of CD4^+^ and CD8^+^ T cells expressing CCR5 in blood from the SIV-infected infant RMs. (A,B) Changes in the CCR5^+^ CD4^+^ T cells; (C,D) Changes in the CCR5^+^ CD8^+^ T cells; Left panels illustrate individual animals; right panels averages. Red symbols illustrate MVC-treated animals. Open black symbols illustrated untreated controls. Vertical bars in the right panels are the standard errors of the means.

### MVC treatment had no discernable impact on the levels of T cell activation and proliferation in SIV-infected infant RMs

These analyses were prompted by studies reporting either that MVC treatment results in a resolution of chronic immune activation that goes beyond the levels of viral control (48) or, conversely, that MVC administration increases the levels of T-cell activation (49). While SIV infection was associated in both MVC treated and untreated RMs with increased levels of CD4^+^ and CD8^+^ T cell proliferation (Figure 10A and B) and immune activation (Figure 10C and D), no significant difference was observed throughout the follow-up between the two groups. We conclude that MVC administration did not significantly influence the levels of CD4^+^ and CD8^+^ T cell immune activation and proliferation in SIV-infected infant RMs.

**Figure 10.**
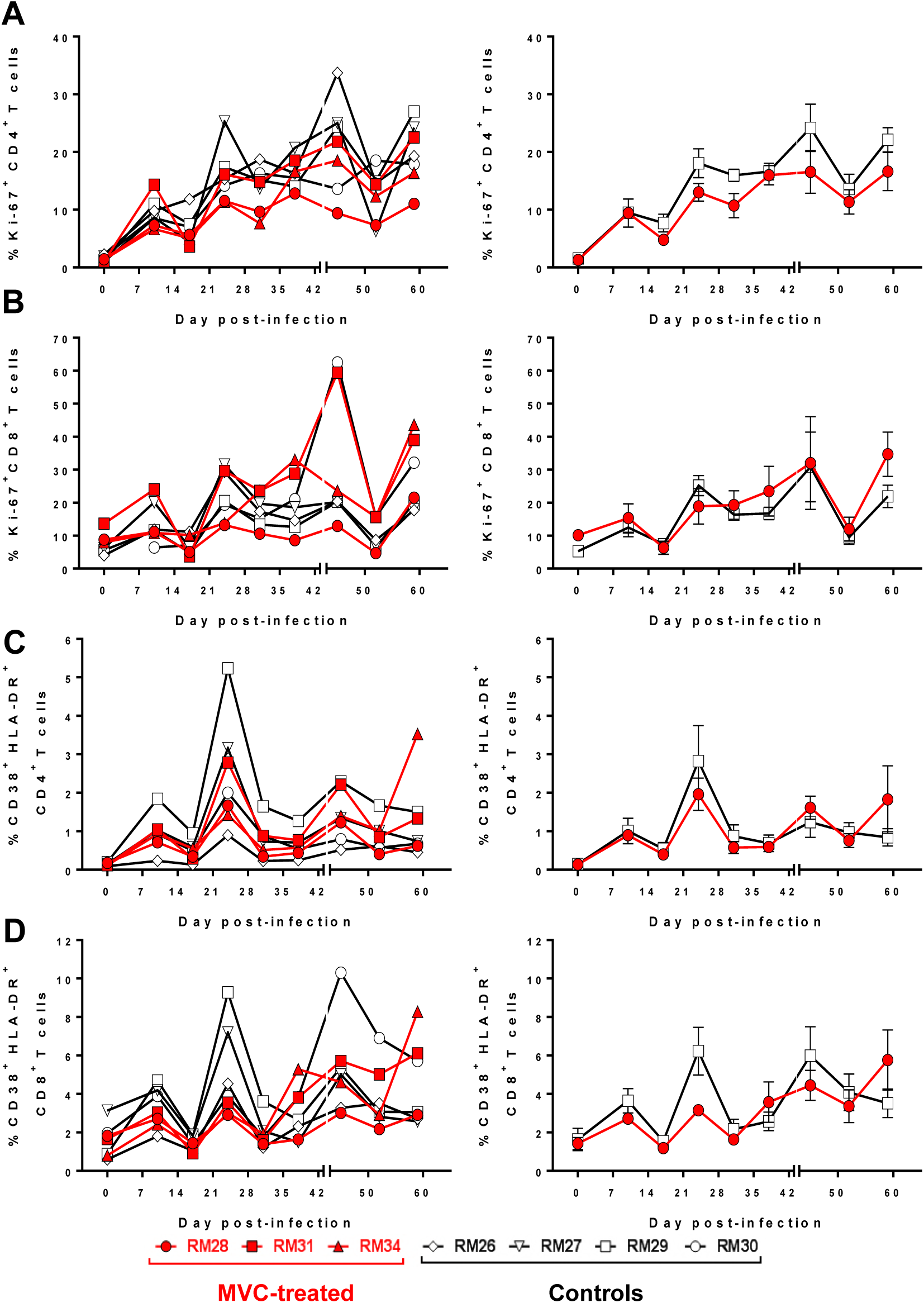
Changes in the frequency of CD4^+^ and CD8^+^ T cells expressing proliferation and immune activation markers in blood from the SIV-infected infant RMs. (A) frequency of the CD4^+^ T cells expressing proliferation marker Ki-67; (B) frequency of the CD8^+^ T cells expressing proliferation marker Ki-67; (C) frequency of the CD4^+^ T cells expressing immune activation markers CD38 and HLA-DR; (D) frequency of the CD8^+^ T cells expressing immune activation markers CD38 and HLA-DR. Left panels illustrate individual animals; right panels averages. Red symbols illustrate MVC-treated animals. Open black symbols illustrated untreated controls. Vertical bars in the right panels are the standard errors of the means. Mann-Whitney test was used to calculate the exact p value.

**Figure 11.**
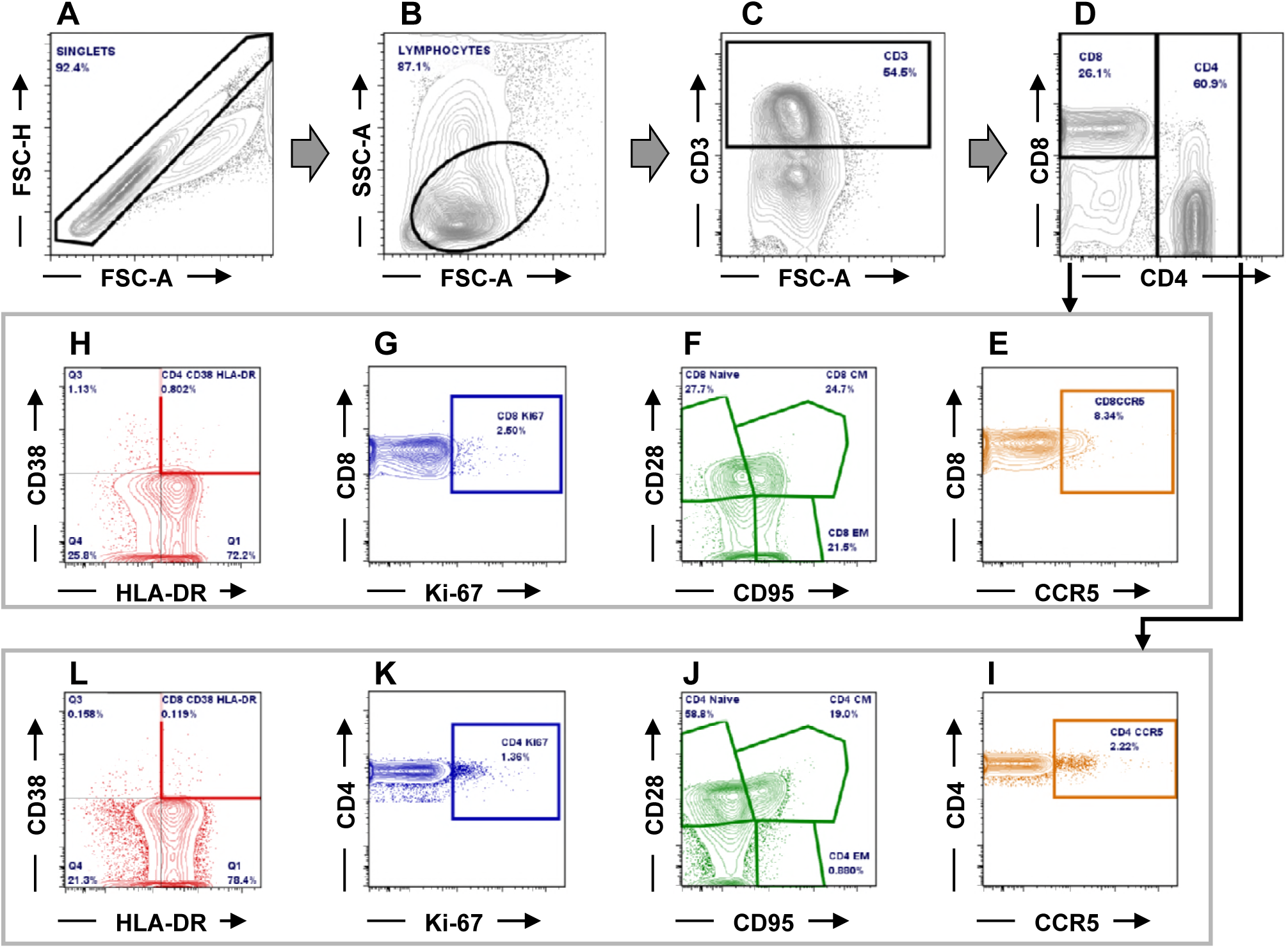
Gating strategy employed to characterize the CD4^+^ and CD8^+^ T cells and their levels of expression for CCR5, as well as the frequency of activated and proliferating T cells (Illustrative plots from RM34) (a-d) CD4^+^ and CD8^+^ T cells were gated on singlets followed by lymphocytes and CD3^+^; (e) CD4^+^ and (i) CD8^+^ T cells expressing CCR5; (f) CD4^+^ and (j) CD8^+^ T-cell naïve and memory subsets; (g) CD4^+^ and (k) CD8^+^ T cells expressing Ki-67; (h) activated CD4^+^ and (l) CD8^+^ T cells expressing CD38 and HLA-DR. FSC-A: forward scatter area; FSC-H: forward scatter height; SSC-A: side scatter area.

## Discussion

While breastfeeding is the healthiest practice for feeding infants, breast milk can also transmit SIV/HIV infection when mothers are infected (50). Without prevention, 13-48% of babies born to HIV-1-infected mothers become HIV infected (4). Perinatal administration of short-term ART to HIV-infected mothers dramatically decreases the rates of HIV-1 MTIT (51). Yet, even with ART prophylaxis, breastfeeding transmission accounts for half of the MTIT cases (52), with overall HIV breastfeeding transmission rates being of approximately 13%, higher in the mothers that seroconvert postpartum (29%) (52) or are in the AIDS stage (37%) (53). Administration of ART to breastfeeding mothers and prolonged ART prophylaxis to infants significantly impacted HIV breastfeeding transmission (54), but this strategy has yet to assess the rates of residual transmission; the degree of drug toxicity on the infant; and the risk for transmission/selection of drug-resistant viruses. Also, to be effective, this strategy has still to circumvent multiple barriers related to implementation (27).

HIV breastfeeding transmission is devastating in developing countries, where 95% of babies are breastfed for up to 2 years (55) and where the benefits of breastfeeding outweigh the risks of breastfeeding transmission (55), as the use of replacement feeding is hindered by access to clean water, cost, availability of formula and cultural background (51). Strategies allowing HIV-infected mothers to breastfeed, while completely controlling breastfeeding transmission, are badly needed.

The rates of SIV MTIT are negligible in African NHP species (AGMs, mandrills or sooty mangabeys) that are natural hosts of SIVs (24-26, 56), in spite of the fact that milk VLs are comparable to those observed in pathogenic infections (57). In experimental studies, we failed to document any SIV breastfeeding transmission in mandrills, even during the acute infection of lactating dams (24). Meanwhile, these low rates of SIV breastfeeding transmission are associated with low levels of mucosal target cells (24, 26), and we documented that, in experimental conditions, susceptibility to mucosal SIV infection of natural hosts is age-related and correlates with the availability of target cells at the mucosal sites (28). These features led us to hypothesize that the mucosal milieu of breastfed infant represent an effective barrier to HIV breastfeeding transmission and that the experimental blockade of mucosal target cell availability may represent an effective new strategy to prevent HIV breastfeeding transmission.

There are multiple rationales for blocking CCR5 to prevent HIV transmission, the most important of which being that CCR5 is the main coreceptor for HIV-1 (58, 59), being thus relevant for the first steps of infection; furthermore, transmitted/founder viruses always use CCR5 for virus entry (60). CCR5 antagonists inhibit replication of R5-tropic HIV variants by blocking viral entry into the target cells (61). MVC is the only FDA-approved CCR5 antagonist (62) and blocks HIV-1 entry by binding CCR5 and suppressing viral infection (63). In addition to modulating CCR5 expression and function (64), MVC may have immunomodulatory effects by blocking binding of the natural ligands of CCR5 (MIP-1α, MIP-1β and RANTES) (65). As a result, CCR5 blockade in HIV-infected subjects reduces immune activation and improve CD4^+^ T cell restoration (66, 67). For the CCR5 blockade, we used MVC, which is FDA-approved, reasoning that, if proven effective, our strategy could be readily implemented to prevent HIV breastfeeding transmission.

Similar to previous studies on MVC safety and tolerance (68, 69), orally administered MVC was safe and well tolerated in all infant RMs, without any obvious side effects, adverse reactions or an impact on the development of the immune system of the infants due to the blockade of a chemokine that may decisively contribute to the immune system maturation (70). As such, we concluded that prolonged CCR5 blockade did not have any deleterious effects on the immune effectors.

Surprisingly, systemic MVC administration only marginally impacted oral SIVmac transmission to infant RMs. At the end of the SIV challenge experiments, when all the RMs in the untreated control group were infected with SIVmac, 60% (3 out of 5 RMs) of the MVC-treated infant RMs were also infected. Furthermore, MVC treatment had no effect on the number of viral challenges needed to transmit SIV or the outcome of SIV infection in infant RM. The only discernible difference observed between the SIV-infected MVC-treated and untreated infant RMs was a significant delay of ramp-up viremia in the MVC-treated infants.

This lack of efficacy of MVC in preventing oral SIV transmission to infant RMs and its limited impact on the key parameters of SIV infection in SIV-infected infant RMs might be due to multiple causes, such as: (i) underdosing of MVC which could limit its therapeutic efficacy; (ii) overexposure to the virus during the challenge experiments, which might have offset the effects of MVC; and (iii) biological promiscuity of SIVs, that may use other coreceptors than CCR5 to infect its target cells (71-73). To examine our MVC dosing strategy, we performed two sets of experiments: first, as the MVC interactions with CCR5 might be different in macaques compared to humans, we sought to demonstrate that MVC successfully blocks CCR5 in infant RMs and to this end we performed an occupancy test (69). In this test, the binding of MVC to CCR5 coreceptors prevents internalization of CCR5 by MIP-1β and thus, the degree of CCR5 internalization by MIP-1β provides an indirect measurement of MVC binding to CCR5. The occupancy test demonstrated that, at the time of viral challenge, 4 hours after oral administration of MVC, CCR5 internalization was robustly blocked. Similarly, testing of the samples collected just prior to drug administration showed that the minimal levels of MVC were generally sufficient to compete with the virus for CCR5 coreceptor occupancy. Note, however, that the lowest coreceptor occupancy was observed in tonsils, a potential site of virus entry upon oral transmission (74).

Next, we measured the MVC plasma concentrations at the time of virus challenge and we documented that steady-state exposure of MVC was similar to therapeutic concentrations in HIV-infected patients. In a different group of infant RMs, we performed MVC dosage in tissues and showed that the drug reaches steady levels in both tonsils and at the mucosal sites, suggesting that the dose of MVC employed here was sufficient to realize a clinical effect. We noted, however, that the minimal levels of MVC, measured just prior to the morning administration of the drug were low and, at least in two instances, below the limits of detection of the assay. Interestingly, the infant RMs which remained uninfected at the completion of the study also had very low minimal levels of MVC, suggesting that the clinical dose used here is probably sufficient for prevention. Nevertheless, drug monitoring revealing a relatively abrupt decline in the MVC levels in infant RMs may also point to a different metabolism of the drug in RMs compared to humans, thus calling for a more detailed evaluation. This aspect is particularly important, as the MVC effect observed here was marginal and, in one case (RM28), a documented undetectable plasma level of MVC was followed by SIV infection.

To rule out virus overdosing, we performed single genome amplification of the molecular tag (32, 75) and showed that none of the MVC-treated or control infant RMs were infected with more than one viral variant, thus discarding the eventuality of an SIVmac overexposure that could have offset the protective effect of MVC.

Finally, to discard the hypothesis of a more promiscuous receptor usage of SIVmac compared to HIV-1, we assessed the coreceptor usage of SIVmac766XII. Differently from HIV-1, which uses CCR5 as the main coreceptor and may expand to use CXCR4 in advanced stages of infection, the majority of SIV strains are more promiscuous, being able to use, in addition to CCR5, BOB/GPR15, (76, 77) and Bonzo/STRL33 (78, 79) for efficient entry into the target cells. More recently, multiple SIV strains were reported to use alternative coreceptors for viral entry, most notably CXCR6 (71-73, 80). This coreceptor usage pathway was reported for the SIVs isolated from both AGMs and sooty mangabeys (71, 73, 80). However, testing of our viral stock for coreceptor usage, clearly demonstrated that CCR5 is the only coreceptor used by SIVmac766XII *in vivo*, similar to the transmitted/founder HIV-1 strains, and validating our choice of challenge strain. While SHIV env strains might have been preferable to SIVmac in this study, fully functional transmitted/founder SHIVs only became available after the completion of this study (81, 82)

As such, our study indicates that MVC was relatively efficient in blocking CCR5 and well tolerated in infant RMs, but exerted only a marginal effect on SIV oral transmission. Failure to block infection was not due to underdosing of MVC, overexposure to the virus during the challenge phase, or alternative coreceptor usage. While a more rapid MVC metabolism in RMs might have impacted MVC efficacy to prevent infection, additional studies would be needed to explore the prophylactic efficacy of target cell blockade for preventing oral HIV transmission through breastfeeding.

Finally, note that systemic administration of MVC did not prevent intrarectal transmission of SHIV (37). Furthermore, CCR5 blockade with MVC was reported to be ineffective in preventing rectal HIV transmission in humans (83). As such, it is possible that CCR5 blockade by MVC is not sufficiently effective in preventing HIV transmission and ineffective in blocking the CCR5 and the use of newly, more potent CCR5 inhibitors will have a better effect in preventing oral SIV transmission, as recently suggested (84).

## Material and Methods

### Ethics statement

Eleven RMs aged six month old were included in this study. They were all housed and maintained at the Plum Borough animal facility of the University of Pittsburgh in agreement with the standards of the Association for Assessment and Accreditation of Laboratory Animal Care (AAALAC). The RMs were fed and housed according to regulations set forth by the Animal Welfare Act and the *Guide for the Care and Use of Laboratory Animals* (85). The RM infants were socially housed indoors in stainless steel cages, had 12/12 light cycle, were fed twice daily, water was provided ad libitum. They were also given various toys and feeding enrichments. The animals were observed twice daily and any signs of disease or discomfort were reported to the veterinary staff for evaluation. For sample collection, animals were anesthetized with 10 mg/kg ketamine hydrochloride (Park-Davis, Morris Plains, NJ, USA) or 0.7 mg/kg tiletamine hydrochloride and zolazepan (Telazol, Fort Dodge Animal Health, Fort Dodge, IA) injected intramuscularly. After viral challenge, all the infant RMs that became infected were followed for 120 days and sacrificed by intravenous administration of barbiturates prior to the onset of any clinical signs of disease. The animal studies were approved by the University of Pittsburgh Institutional Animal Care and Use Committee (IACUC) (Protocol #13112685).

### Virus stock

The SIVmac766XII stock (Figure 1) is composed of parental SIVmac766, previously identified as a transmitted/founder virus and infectious molecular clone (38) and eleven other viral variants differing from the parental clone by 3 synonymous mutations in integrase similar to the SIVmac239X swarm previously described (32). The virus stock was generated by transfecting 293T cells with equal amounts of each of the 12 molecularly modified plasmids for 24 h using the TransIT HEK-293 transfection reagent (Mirus Bio) according to the manufacturer’s instructions. Culture medium was changed at 24 h post-transfection and again at 48 h posttransfection. At 72 h posttransfection, virus-containing supernatant was clarified by centrifugation, sterile-filtered through a 0.45 µM filter, aliquoted, and stored at −80°C. A series of small scale cotransfections with subsequent sequencing analyses to determine the relative proportion of each tagged variant in the virus pool was used to identify the relative input proportion of each plasmid in the DNA cotransfection pool that would yield roughly equal proportions of tagged viruses in the SIVmac766XII stock. Virus titers were determined using TZM-bl reporter cells (NIH AIDS Research and Reference Reagent Program), which contain a Tat-inducible luciferase and a β-galactosidase gene expression cassette. Infectious titers were measured by counting individual β-galactosidase-expressing cells per well in cultures infected with serial 3-fold dilutions of virus. Wells containing dilution-corrected blue cell counts within the linear range of the virus dilution series were averaged to generate an infectious titer in infectious units (IU) per milliliter. The SIVmac766XII stock contained 2.75×10^5^ IU/ml.

### MVC treatment and animal infection

Five RMs received a total daily dose of 300 mg/kg administered as two divided doses (150 mg/kg) per os with food. The MVC dose was allometrically scaled to twice the dose of humans (300 mg/kg). One month after MVC initiation, all the MVC-treated infants, together with 4 untreated infant RMs were orally administered 10,000 IU of SIVmac766XII. Virus challenge occurred 4 hours after the morning MVC administration, when the concentrations of MVC were demonstrated to be maximal. Viral challenge was repeated every two weeks, for up to 6 times. At the time of viral challenges, CCR5 coreceptor occupancy (33) was also closely monitored. To evaluate the concentration and pharmacokinetics of the MVC in the tissues, we enclose in our experimental design two RMs treated with MVC similarly to the infants in the study group, but not challenged.

### Sampling

At the time of virus challenge, blood (1.5 ml) was collected into EDTA-CPTs, to monitor coreceptor occupancy and MVC plasma levels, and then every three days, to monitor SIV infection. Once an animal was demonstrated to be SIV infected, sampling was scheduled to monitor the acute and early chronic infection (10, 17, 24, 31, 38, 45, 59 dpi). Superficial LNs, tonsils and intestine were collected just from the two MVC-treated SIV-unexposed infant RM controls. To prevent changes in CCR5 expression due to storage and shipping of unprocessed peripheral blood mononuclear cells, all blood and tissue samples were processed within 60 min from collection.

### Analysis of plasma MVC concentration

MVC concentrations were measured in plasma samples collected 4 hours after administration of one oral dose of 150 mg/kg by a validated liquid chromatography–tandem mass spectrometry (LC-MS/MS) method using a Shimadzu high-performance liquid chromatography system for separation, and an AB SCIEX API 5000 mass spectrometer (AB SCIEX, Foster City, CA, USA) equipped with a turbo spray interface for detection. The calibrated linear range was 5-5000 ng/ml in plasma. All samples were extracted by protein precipitation with a stable, isotopically-labeled internal standard (MVC-d6) added for quantification. All calibration standards and quality controls (QCs) were prepared in drug-free NHP plasma. Calibration standards and QC samples met 15% acceptance criteria for precision and accuracy. Plasma MVC concentrations were expressed as ng of MVC per ml.

### Analysis of tissue MVC concentration

MVC concentrations in LNs, tonsils and intestine were measured on samples collected either 4 hours after administration of an oral dose of 150 mg/kg MVC, or immediately before MVC administration. Tissue MVC concentrations were measured with the same methodology used to measure plasma MVC concentrations and were expressed as ng of MVC per gr of tissue (ng/gr of tissue).

### MIP-1 internalization assay

The coreceptor occupancy test was performed to assess MVC binding to CCR5 in blood and in LNs, tonsils and intestine (33, 69, 86). The principle of this test is that the binding of MVC prevents internalization of CCR5 by MIP-1β and thus, the degree of CCR5 internalization by MIP-1β provides an indirect measurement of MVC binding to CCR5.

PBMC-rich plasma samples were isolated by centrifugation of CTP tube at 2,200 rpm × 25 minute. The cell pellets were resuspended in the plasma at 5 × 10^6^ cell/ml obtaining cell-enriched plasma samples. For the assay in blood, 5 × 10^5^ cell-enriched plasma sample (100 µl) was aliquoted into three separately labeled 5 ml Facs tubes containing the isotype control (Tube 1), the MVC-stabilized CCR5 (Tube 2) and the test sample (Tube 3). Cells from LNs, tonsils and intestinal biopsies were collected as described (87) and 5 × 10^6^ cells were resuspended in the plasma and aliquoted (100 µl) in three tubes as described above for blood. Fifty µl of MVC 1µM (CCR5 stabilizing solution) were added to Tube 2; the same volumes of 50 µl of 1% PBS/BSA (w/v) were added to Tube 1 and 3. Tubes 1-3 were briefly vortexed and incubated at 37ºC for 30 min, followed by centrifugation at 1500 rpm for 5 min. The supernatant was discarded, and 15 µl of MIP-1β (100 nM) was added to all tubes. The mixture was then vortexed and incubated uncapped for 45 min at 37ºC to enable CCR5 internalization. Then, one ml of 0.5% paraformaldehyde in PBS was added to each tube, which were then vortexed and incubated in the dark at room temperature (RT) for 10 min. Cells were then washed with PBS by centrifugation at 1500 rpm for 5 min, and stained with a combination of antibodies (Table 1) appropriate for the identification of CD4^+^ and CD8^+^ T cells expressing CCR5, and a combination of isotype and fluorescence-minus-one controls. Labeled cells were washed once with 1% FBS PBS, fixed in 2% formaldehyde PBS (Affimetrix, Santa Clara, CA) and then acquired the same day on a custom four-laser BD LSRII instrument (BD Bioscience). Only singlet events were gated and a minimum of 250,000 live CD3 cells were acquired. Populations were analyzed using FlowJo software version 7.6.5 (Tree Star Inc. Ashland, OR) and the graphs were generated with GraphPad Prism 6.04. The percentage of receptor occupancy was calculated using CCR5 expression data obtained for PBL aliquots incubated with chemokine in the presence of 1 µM MVC (Tube 2) and in the absence of additional MVC (Tube 3), as follows: % receptor occupancy = (% of CCR5 expression in Tube 3)/(% of CCR5 expression in Tube 2)×100.

### Alternative coreceptor usage by SIVmac76XII in CF2th-Luc cells

CF2th-Luc cells, which contain a Tat-driven luciferase reporter, were transfected using Fugene 6 (Promega) with two expression plasmids: one containing RM CD4 and one containing coreceptor (or pcDNA3.1 empty vector). Cells were infected 48 hours later with SIVmac766XII (using 2,750, 13,750 or 27,500 IU) by spinoculation for 2 hours at 1,200 × *g*. Cells were incubated for 48 hours at 37°C, then they were lysed and luciferase content quantified as relative light units (RLU), as previously described (71).

### Alternative coreceptor usage by SIVmac76XII in PBMCs

Cryopreserved PBMC from two RMs (RM1 and RM2) were thawed in complete medium (RPMI 1640 with 10% FBS, 1% L-glutamine and 1% penicillin/streptomycin) and stimulated for three days with 5 µg/ml Concanavalin A and 100 U/ml IL-2. Activated PBMCs were plated at 2×10^5^ per well in 96-well plates, pretreated for one hour with 15 µM MVC (NIH AIDS Reagent Program) or with vehicle alone (DMSO), and then infected with SIVmac766XII (550 IU) by spinoculation for 1.5 hours at 1,200 g. Cells were then washed, incubated at 37°C, and supernatants were collected and 50% media replaced every 3-4 days. Replication was measured by SIV p27 Gag antigen production in the supernatant by ELISA (Perkin-Elmer).

### Assessment of the T/F virus variants

The number of T/F variants was determined using a real-time single genome amplification (RT-SGA) approach described previously (32). Briefly, a 300 bp portion of the integrase gene surrounding the mutated site was amplified and sequenced using a limiting dilution PCR where only a single genome is amplified (SGA) per reaction. Viral RNA was extracted using the QIAamp Viral RNA Mini Kit (Qiagen) and then reverse transcribed into cDNA using SuperScript III reverse transcription according to manufacturer’s recommendations (Invitrogen) and the antisense primer SIVmacIntR1 5’-AAG CAA GGG AAA TAA GTG CTA TGC AGT AA-3’. PCR was then performed with 1 × PCR buffer, 2 mM MgCl_2_, 0.2 mM of each deoxynucleoside triphosphate, 0.2 μM of each primer, and 0.025 U/μl Platinum Taq polymerase (Invitrogen) in a 10-μl reaction with sense primer SIVmacIntF1 5’-GAA GGG GAG GAA TAG GGG ATA TG-3’ and antisense primer SIVmacIntR3 5’-CAC CTC TCT AGC CTC TCC GGT ATC C-3’ under the following conditions: 1 cycle of 94°C for 2 min, 40 cycles at 94°C for 15 sec, 55°C for 30 sec, 60°C 1.5 min, and 72°C for 30 sec. Template positive reactions were determined by real-time PCR using as gene specific probe SIVIntP 5’-TCC CTA CCT TTA AGA TGA CTG CTC CTT CCC CT-3’ with FAM6 and ZEN/Iowa Black Hole Quencher (Integrated DNA Technologies) and directly sequenced with SIVmacIntR3 using BigDye Terminator technology (Life Technologies). To confirm PCR amplification from a single template, chromatograms were manually examined for multiple peaks, indicative of the presence of amplicons resulting from PCR-generated recombination events, Taq polymerase errors or multiple variant templates.

### Flow cytometry

Flow cytometry was used to assess changes in CD4^+^ and CD8^+^ T cell populations, the frequency of CD4^+^ and CD8^+^ cells expressing CCR5, as well as their proliferation and immune activation status, as described (88-90). Briefly, 2 × 10^6^ cells were stained with viability dye (Blue dye, Life Technologies) and incubated for 15 min in the dark at RT. The cells were than washed with PBS and stained for 30 min at RT in the dark with combinations of antibodies and combinations of isotype and fluorescence-minus-one controls (Table 1) appropriate for the identification of different T cell populations (Figure 111). Stained cells were washed in 1x PBS, fixed with 2% paraformaldheyde solution (PFA) and stored at 4º C prior to acquisition. For intracellular staining, viable cells stained as described above were washed with 1x PBS, permeabilized with a solution containing 0.1% of saponine and incubated for 30 min at room temperature in the dark. Cells were then stained with an anti-Ki-67 antibody (Table 1). Cells were then washed with 1x PBS, fixed with 2% PFA and stored at 4º C prior to the acquisition. A minimum of 250,000 CD3 live cells were acquired with FACSDiva software v.8.0. Acquired cells were analyzed using FlowJo 7.6.5 software.

### Statistical analyses

All statistical analyses were performed with GraphPad Prism Software v.6.02 (GraphPad Software Inc. San Diego CA, USA). Data were expressed as averages ± standard error of means (SEM). Unpaired nonparametric Mann-Whitney t test was used for significant differences between the experimental groups, with regards to the absolute cell counts and frequency of cells expressing CCR5, immune activation and cell proliferation markers. Wilcoxon paired non-parametric test was used, to determine significant differences between the baselines of mean cell frequencies and absolute counts and single time point of MVC treatment, for each experimental group. Chi-square test was used to establish significant differences between the animals that became infected in both groups. Differences were considered statistically significant at p ≤ 0.05.

## Author contribution

EBC, CA and IP designed experiments; EBC, CA, IP, ADK, RGC, BFK analyzed data, BFK and KM provided virus stock and sequence analysis; EBC, CX, KSW, MLC, BBP, KDR, TD, GSHR, DM, and KM performed experiments; EBC, CA, IP BFK, MLC, ADK, KSW, and RGC wrote the manuscript.

## Acknowledgements

We would like to thank Drs. Guido Silvestri and Isaac Rodriguez-Chavez for helpful discussion. This work was supported by grant R56 DE023508 (CA) from the National Institute of Dental and Craniofacial Research (NIDCR). Parts of this study were supported with funds from grants: RO1 HL117715 (IP), R01 AI119346 (CA) and R01 HL123096 (IP). KDR and BBP were supported by T32 AI065380 (Pitt AIDS Research Training, PART). BBP was supported by RO1 AI104373. This work was also partially supported by the UNC Center for AIDS Research (CFAR) (P30 AI50410). Finally, this project has been funded in part with Federal funds from the National Cancer Institute, National Institutes of Health, under Contract No. HHSN261200800001E. The funders had no role in study design, data collection and analysis, decision to publish, or preparation of the manuscript. The content of this publication does not necessarily reflect the views or policies of the Department of Health and Human Services, nor does mention of trade names, commercial products, or organizations imply endorsement by the U.S. Government.

